# SARS-CoV-2 spike P681R mutation, a hallmark of the Delta variant, enhances viral fusogenicity and pathogenicity

**DOI:** 10.1101/2021.06.17.448820

**Authors:** Akatsuki Saito, Takashi Irie, Rigel Suzuki, Tadashi Maemura, Hesham Nasser, Keiya Uriu, Yusuke Kosugi, Kotaro Shirakawa, Kenji Sadamasu, Izumi Kimura, Jumpei Ito, Jiaqi Wu, Kiyoko Iwatsuki-Horimoto, Mutsumi Ito, Seiya Yamayoshi, Seiya Ozono, Erika P Butlertanaka, Yuri L Tanaka, Ryo Shimizu, Kenta Shimizu, Kumiko Yoshimatsu, Ryoko Kawabata, Takemasa Sakaguchi, Kenzo Tokunaga, Isao Yoshida, Hiroyuki Asakura, Mami Nagashima, Yasuhiro Kazuma, Ryosuke Nomura, Yoshihito Horisawa, Kazuhisa Yoshimura, Akifumi Takaori-Kondo, Masaki Imai, The Genotype to Phenotype Japan (G2P-Japan) Consortium, So Nakagawa, Terumasa Ikeda, Takasuke Fukuhara, Yoshihiro Kawaoka, Kei Sato

## Abstract

During the current SARS-CoV-2 pandemic, a variety of mutations have been accumulated in the viral genome, and currently, four variants of concerns (VOCs) are considered as the hazardous SARS-CoV-2 variants to the human society^1^. The newly emerging VOC, the B.1.617.2/Delta variant, closely associates with a huge COVID-19 surge in India in Spring 2021^2^. However, its virological property remains unclear. Here, we show that the B.1.617.2/Delta variant is highly fusogenic, and notably, more pathogenic than prototypic SARS-CoV-2 in infected hamsters. The P681R mutation in the spike protein, which is highly conserved in this lineage, facilitates the spike protein cleavage and enhances viral fusogenicity. Moreover, we demonstrate that the P681R-bearing virus exhibits higher pathogenicity than the parental virus. Our data suggest that the P681R mutation is a hallmark that characterizes the virological phenotype of the B.1.617.2/Delta variant and is closely associated with enhanced pathogenicity.

## Main

In December 2019, an unusual infectious disease, now called COVID-19, emerged in Wuhan, Hubei province, China^3, 4^. SARS-CoV-2, the causative agent of COVID-19, has rapidly spread all over the world, and as of July 2021, SARS-CoV-2 is an ongoing pandemic: more than 180 million cases of infections have been reported worldwide, and more than 4 million people died of COVID-19^1^.

During the current pandemic, SARS-CoV-2 has acquired a variety of mutations^5^. First, in the spring of 2020, a SARS-CoV-2 derivative harbouring the D614G mutation in its spike (S) protein has emerged and quickly become predominant^6^. Because the D614G mutation increases viral infectivity, fitness, and inter-individual transmissibility^7–12^, the D614G-bearing variant has quickly swept out the original strain. Since the fall of 2020, some SARS-CoV-2 variants bearing multiple mutations have emerged and rapidly spread worldwide. As of June 2021, there have been at least five variants of concern (VOC): B.1.1.7 (Alpha), B.1.351 (Beta), P.1 (Gamma), B.1.427/429 (Epsilon; note that this variant is currently out of concerns/interest) and B.1.617.2 (Delta), and these lineages respectively emerged in the UK, South Africa, Brazil, the USA and India^13, 14^.

At the end of 2020, the B.1.617 lineage has emerged in India, and this variant is thought to be a main driver of a massive COVID-19 surge in India, which has peaked 400,000 infection cases per day^2^. The B.1.617 lineage includes three sublineages, B.1.617.1, B.1.617.2 and B.1.617.3, and a sublineage, B.1.617.2, is the latest VOC, the Delta variant^13, 14^. Importantly, early evidence have suggested that the B.1.617.2/Delta may have an increased risk of hospitalization compared to the B.1.1.7 cases^15, 16^. However, the virological features of this newly emerging VOC, particularly its infectivity and pathogenicity, remain unclear. Additionally, although recent studies have shown that the B.1.617.2/Delta variant is relatively resistant to the neutralising antibodies (NAbs) elicited by vaccination^17, 18^, the mutation(s) that are responsible for the virological features of this VOC are unaddressed. In this study, we demonstrate that the B.1.617.2/Delta is more pathogenic than the prototypic SARS-CoV-2 in a Syrian hamster model. We also show that the P681R mutation in the S protein is a hallmark mutation of this lineage. The P681R mutation enhances the cleavage of SARS-CoV-2 S protein and enhances viral fusogenicity. Moreover, we demonstrate that the P681R mutation is responsible for the higher pathogenicity of the B.1.617.2/Delta variant *in vivo*.

### Phylogenetic and epidemic dynamics of the B.1.617 lineage

We set out to investigate the phylogenetic relationship of the three subvariants belonging to the B.1.617 lineage. We downloaded 1,761,037 SARS-CoV-2 genomes and information data from the Global Initiative on Sharing All Influenza Data (GISAID) database (https://www.gisaid.org; as of May 31, 2021). As expected, each of three sublineages, B.1.617.1, B.1.617.2 and B.1.617.3, formed a monophyletic cluster, respectively (Fig. 1a, Extended Data Fig. 1). We then analyzed the epidemic of each of three B.1.617 sublineages. The B.1.617 variant, particularly B.1.617.1, was first detected in India on December 1, 2020 (GISAID ID: EPI_ISL_1372093) (Fig. 1b-d). Note that a SARS-CoV-2 variant (GISAID ID: EPI_ISL_2220643) isolated in Texas, the USA, on August 10, 2020, was also recorded to belong to the B.1.617.1. However, the S protein of this viral sequence (GISAID ID: EPI_ISL_2220643) possesses neither L452R nor P681R mutations, both of which are the features of the B.1.617 lineage. Therefore, the EPI_ISL_2220643 sequence isolated in the USA may not be the ancestor of the current B.1.617.1 lineage, and the EPI_ISL_1372093 sequence obtained in India would be the oldest B.1.617 lineage.

**Fig. 1.**
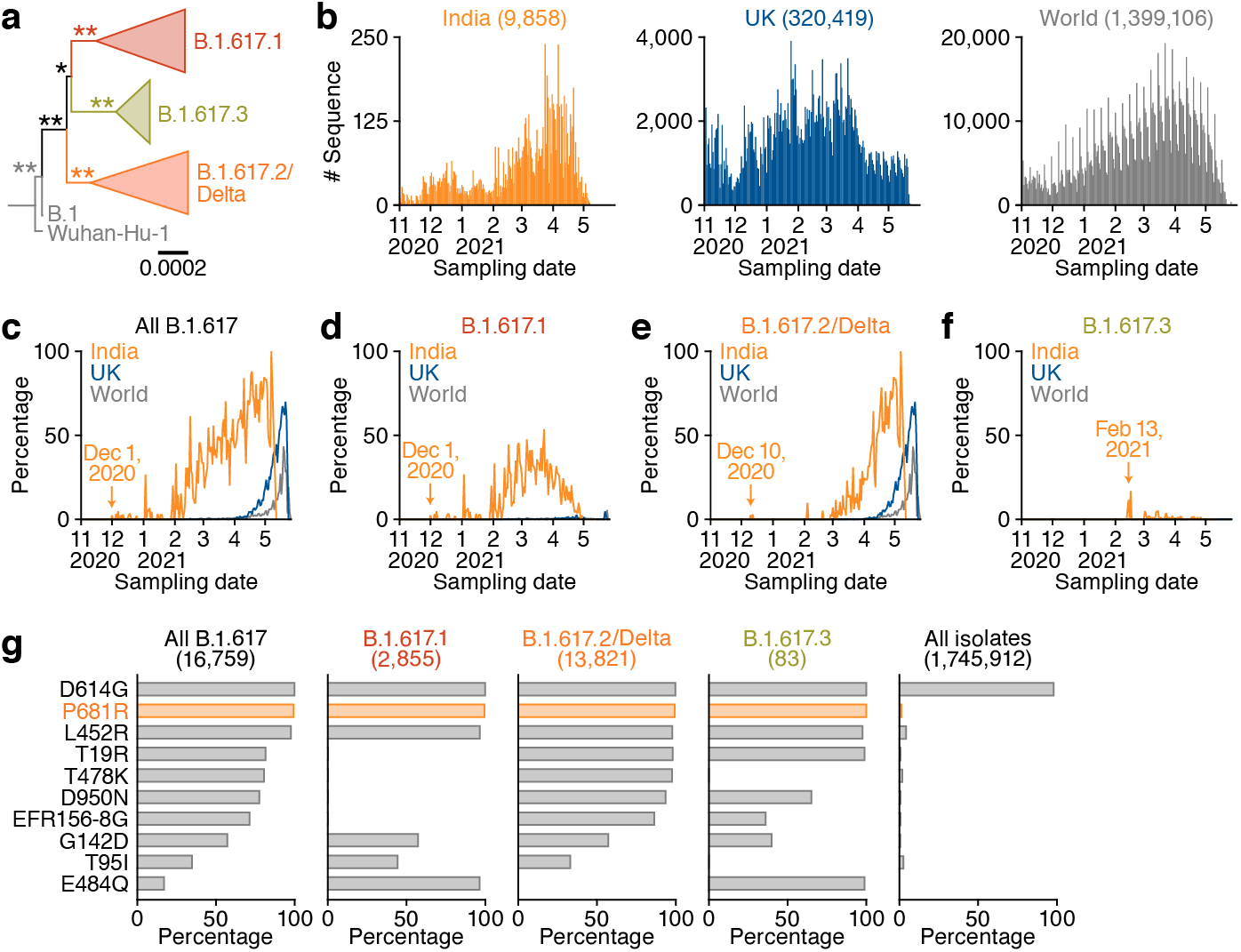
Molecular phylogenetic and epidemic dynamics of the B.1.617 lineage during the pandemic. **a,** A phylogenetic tree of the B.1.617 lineage. Bar, 0.0002 substitutions per site. Bootstrap values, **, 100%; *, >70%. The uncollapsed tree is shown in Extended Data Fig. 1. **b-f,** Epidemic dynamics of the B.1.617 lineage. **b,** The numbers of sequences deposited in GISAID per day for India (orange, left), UK (blue, middle), and the whole world (gray, right). **c-f,** The percentages of each lineage deposited per day (**c**, all B.1.617; **d**, B.1.617.1; **e**, B.1.617.2/Delta; **f**, B.1.617.3) from India (orange), the UK (blue) and the whole world (gray) are shown. The date first identified is indicated. The raw data are summarized in **Extended Data Table 1**. **g,** Proportion of amino acid replacements in the B.1.617 lineage. The top 10 replacements conserved in the S protein of the B.1.617 and its sublineages are summarized. The number in parenthesis indicates the number of sequences included in each panel. The raw data are summarized in **Extended Data Table 2**.

The B.1.617.2 (GISAID ID: EPI_ISL_2131509) and B.1.617.3 (GISAID IDs: EPI_ISL_1703672, EPI_ISL_1703659, EPI_ISL_1704392) were detected in India on December 10, 2020 and February 13, 2021, respectively (Fig. 1e, f). The B.1.617.1 sublineage has peaked during February to April, 2021, in India, and then decreased (Fig. 1d). Although the B.1.617.3 variant has sporadically detected in India (Fig. 1f), the B.1.617.2/Delta lineage has become dominant in India since March 2021 and spread all over the world (Fig. 1e). At the end of May 2021, 100%, 70% and 43.3% of the deposited sequences in GISAID per day from India (May 7), the UK (May 21) and the whole world (May 19) have been occupied by the B.1.617.2 sublineage (Fig. 1e **and Extended Data Table 1**).

We next investigated the proportion of amino acid replacements in the S protein of each B.1.617 sublineage comparing with the reference strain (Wuhan-Hu-1; GenBank accession no. NC_045512.2). As shown in Fig. 1g, the L452R and P681R mutations were highly conserved in the B.1.617 lineage, and notably, the P681R mutation (16,650/16,759 sequences, 99.3%) was the most representative mutation in this lineage. These data suggest that the P681R mutation is a hallmark of the B.1.617 lineage.

### Prominent syncytia formation by the B.1.617.2/Delta variant

To investigate the virological characteristics of the B.1.617.2/Delta variant, we conducted virological experiments using a viral isolate of B.1.617.2 (GISAID ID: EPI_ISL_2378732) as well as a D614G-bearing B.1.1 isolate (GISAID ID: EPI_ISL_479681) in Japan. In Vero cells, the growth of the B.1.617.2/Delta variant was significantly lower than that of the B.1.1 isolate (Fig. 2a). Particularly, the levels of viral RNA of the B.1.617.2/Delta variant at 48 hours postinfection (hpi) was more than 150-fold lower than that of the B.1.1 isolate (Fig. 2a). On the other hand, although the growth kinetics of these viruses was relatively comparable in VeroE6/TMPRSS2 cells and Calu-3 cells (Fig. 2a), microscopic observations showed that the B.1.617.2/Delta formed larger syncytia compared to the B.1.1 virus (Fig. 2b). By measuring the size of the floating syncytia in the infected VeroE6/TMPRSS2 culture, the syncytia formed by the B.1.617.2/Delta infection were significantly (2.7-fold) larger than that by the B.1.1 infection (Fig. 2b). Immunofluorescence assay further showed that the B.1.617.2/Delta-infected VeroE6/TMPRSS2 cells exhibit larger multinuclear syncytia compared to the B.1.1 isolate (Extended Data Fig. 2). These results suggest that the B.1.617.2/Delta variant is feasible for forming syncytia compared to the D614G-bearing B.1.1 virus.

**Fig. 2.**
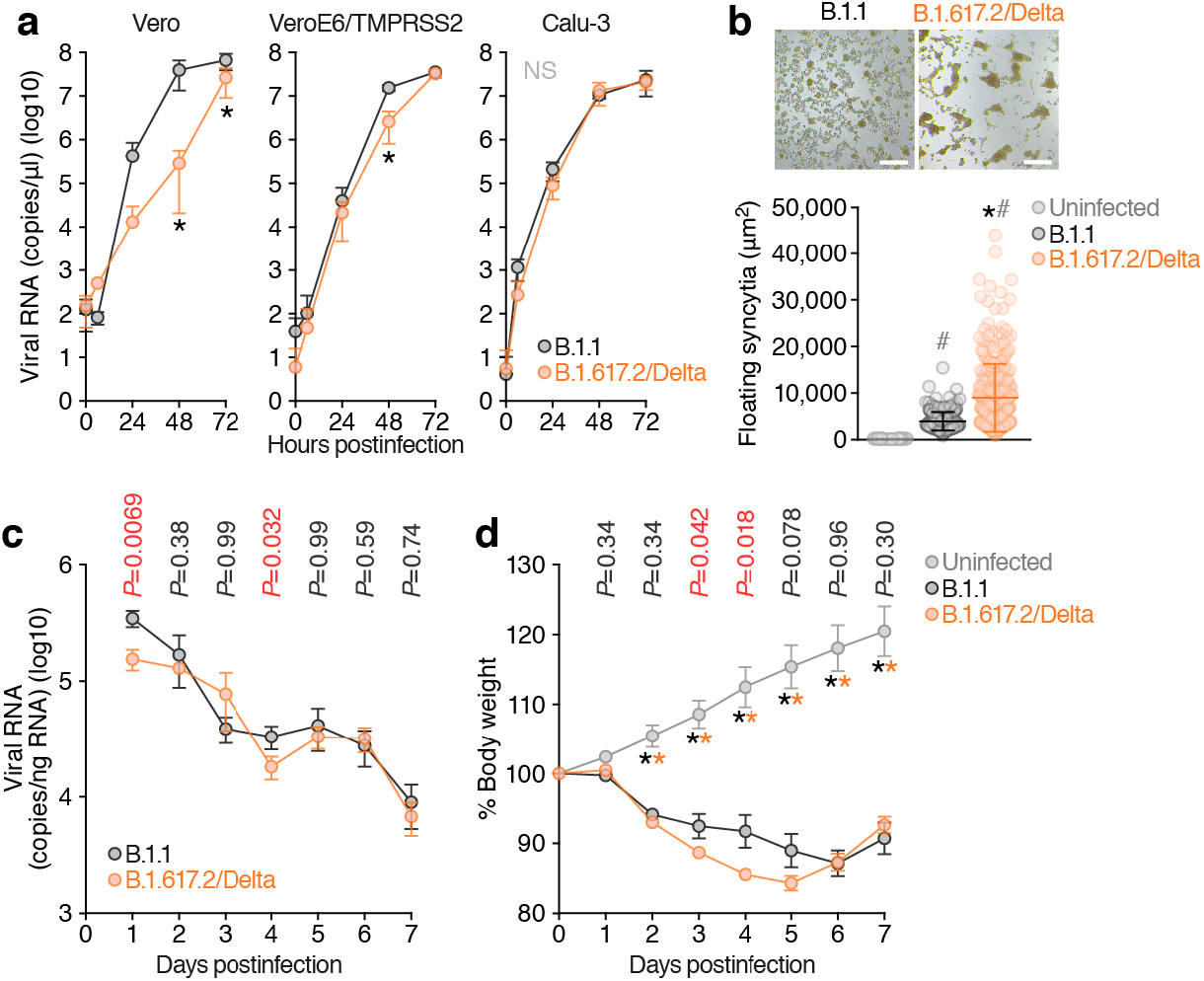
Virological features of the B.1.617.2/Delta variant *in vitro* and *in vivo*. **a,** Growth kinetics of the B.1.617.2/Delta variant and a B.1.1 isolate. A viral isolate of B.1.617.2/Delta (GISAID ID: EPI_ISL_2378732) and a D614G-bearing B.1.1 isolate (GISAID ID: EPI_ISL_479681) [100 50% tissue culture infectious dose (TCID_50_) for Vero cells and VeroE6/TMPRSS2 cells, 1,000 TCID_50_ for Calu-3 cells] were inoculated and the copy number of viral RNA in the culture supernatant was quantified by real-time RT-PCR. The growth curves of the viruses inoculated are shown. Assays were performed in quadruplicate. **b,** Syncytia formation. The syncytia in infected VeroE6/TMPRSS2 cells were observed at 72 hpi. (Top) Representative bright-field images of VeroE6/TMPRSS2 cells at 72 hpi are shown. Bars, 100 μm. (Bottom) The size of floating syncytia in B.1.1-infected (n = 217) and B.1.617.2/Delta-infected (n = 217) cultures are shown. The size of the floating single cells in uninfected culture (n = 177) was also shown as a negative control. **c, d,** Syrian hamster infection with the B.1.617.2/Delta variant. Male hamsters were infected with 10^5^ TCID_50_ of the B.1.1 isolate (n = 6) and the B.1.617.2/Delta isolate (n = 12). Four hamsters at the same age were used for mock infection. The amount of viral RNA in the oral swab (**c**) and body weight (**d**) and were routinely measured. In **a**, statistically significant differences (*, *P* < 0.05) versus the B.1.1 isolate were determined by Student’s *t* test. NS, no statistical significance. In **b**, statistically significant differences versus the B.1.1-infected culture (*, *P* < 0.05) and uninfected culture (#, *P* < 0.05) were determined by the Mann-Whitney U test. In **c** and **d**, statistically significant differences were determined by the Mann-Whitney U test, and those versus uninfected hamsters (*, *P* < 0.05) are indicated by asterisks. The *P* value between the B.1.1 and the B.1.617.2/Delta at each dpi is indicated in the figure.

### Higher pathogenicity of the B.1.617.2/Delta variant in Syrian hamsters

To investigate the pathogenicity of the B.1.617.2/Delta variant, we conducted hamster infection experiments using the B.1.617.2/Delta isolate and the B.1.1 isolate. Although the viral RNA loads in the oral swab of the B.1.617.2/Delta-infected hamsters were significantly lower than those of the B.1.1-infected hamsters at 1 and 4 days postinfection (dpi), these values were comparable any other dpi (Fig. 2c). After infection with these viruses, infected hamsters significantly lost their body weights from 2 dpi. The peak weight loss was 16% for the B.1.617.2/Delta and 13% for the B.1.1, with the B.1.617.2/Delta isolate having a significantly greater weight loss than the B.1.1 at 3 and 4 dpi (Fig. 2d). These results suggest that the B.1.617.2/Delta has a higher pathogenicity than the B.1.1 isolate, despite relatively comparable proliferative potential.

### P681R mutation as the determinant of enhanced and accelerated fusogenicity

The P681R mutation in the S protein is a unique feature of the B.1.617 lineage including the B.1.617.2/Delta variant (Fig. 1g). Because the P681R mutation is located in the proximity of the furin cleavage site (FCS; residues RRAR positioned between 682-5) of the SARS-CoV-2 S protein^19^, we hypothesized that the P681R mutation is responsible for the preference of cell-cell fusion, which leads to larger syncytia formation. To address this possibility, we generated the P681R-bearing artificial virus by reverse genetics (Extended Data Fig. 3) and preformed virological experiments. Although the amounts of viral RNA in the culture supernatants of the D614G/P681R-infected Vero and VeroE6/TMPRSS2 cells were significantly lower than those of the D614G-infected cells in some time points, the growth of these two viruses was relatively comparable (Fig. 3a). However, the size of floating syncytia in the D614G/P681R-infected VeroE6/TMPRSS2 cells at 72 hpi was significantly larger than that in the D614G mutant-infected cells (Fig. 3b). This observation well corresponds to that in the culture infected with the B.1.617.2/Delta variant (Fig. 2b).

**Fig. 3.**
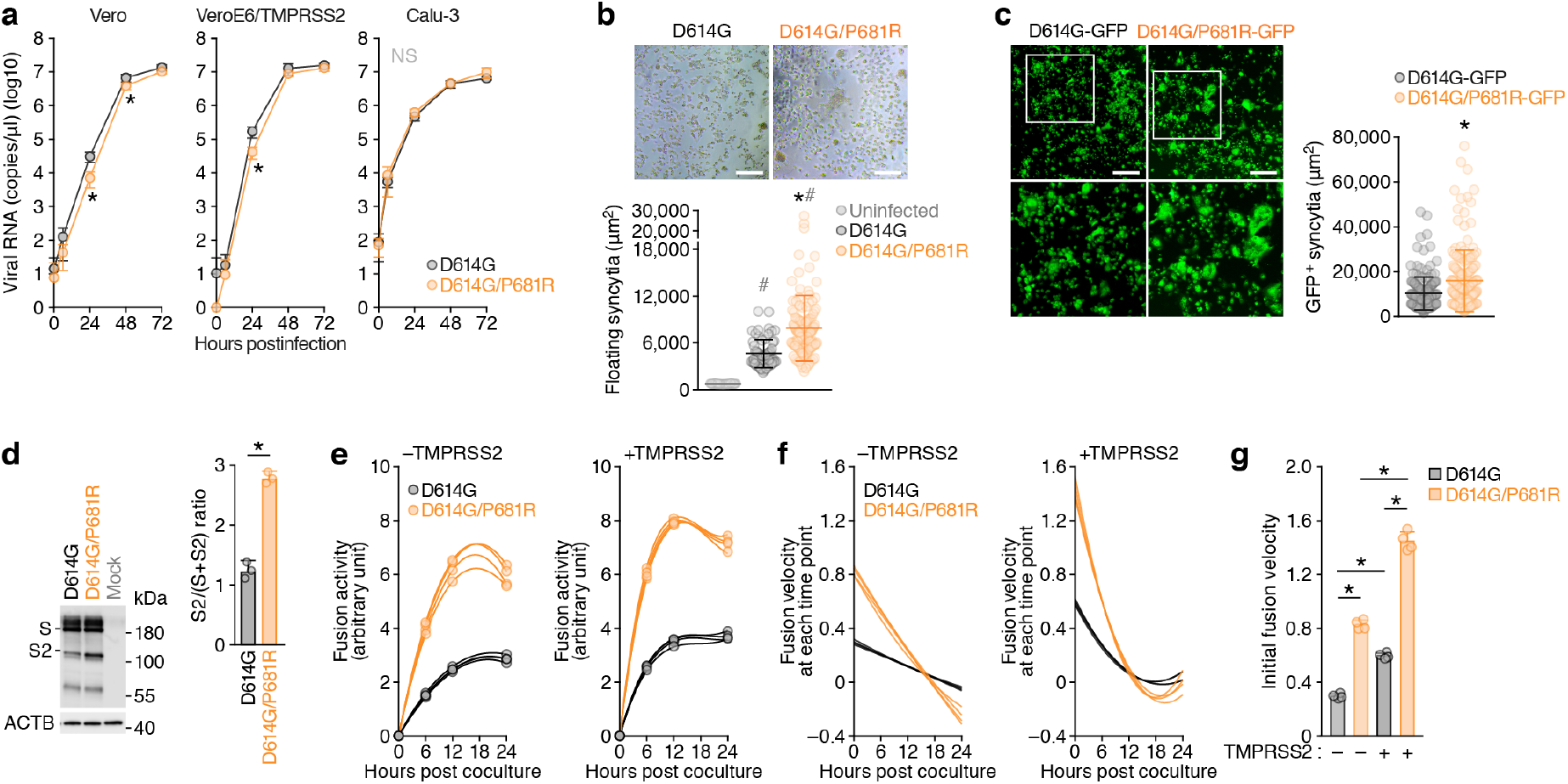
Virological features of the P681R-harbouring virus *in vitro*. **a,** Growth kinetics of artificially generated viruses. The D614G and D614G/P681R mutant viruses were generated by reverse genetics. These viruses (100 TCID_50_) were inoculated into Vero cells and VeroE6/TMPRSS2 cells, and the copy number of viral RNA in the culture supernatant was quantified by real-time RT-PCR. The growth curves of the viruses inoculated are shown. Assays were performed in quadruplicate. **b, c,** Syncytia formation. **b,** (Top) Floating syncytia in VeroE6/TMPRSS2 cells infected with the D614G and D614G/P681R mutant viruses at 72 hpi are shown. Bars, 200 μm. (Bottom) The size of floating syncytia in the D614G mutant-infected (n = 63) and the D614G/P681R mutant-infected (n = 126) cultures are shown. **c,** (Left) Adherent syncytia in VeroE6/TMPRSS2 cells infected with the GFP-expressing D614G and D614G/P681R mutant viruses at 24 hpi are shown. Areas enclosed with squares are enlarged in the bottom panels. Bars, 200 μm. (Right) The size of adherent GFP^+^ syncytia in the D614G mutant-infected (n = 111) and the D614G/P681R mutant-infected (n = 126) cultures are shown. **d,** Western blotting of the S-expressing cells. (Left) Representative blots of SARS-CoV-2 full-length S and cleaved S2 proteins as well as ACTB as an internal control. kDa, kilodalton. (Right) The ratio of S2 to the full-length S plus S2 proteins in the S-expressing cells. **e-g,** SARS-CoV-2 S-based fusion assay. Effector cells (S-expressing cells) and target cells (ACE2-expressing cells or ACE2/TMPRSS2-expressing cells) were prepared, and the fusion activity was measured as described in **Methods**. **e,** Kinetics of fusion activity (experimental data). Assays were performed in quadruplicate, and fusion activity (arbitrary unit) is shown. **f,** The kinetics of fusion velocity estimated by a mathematical model based on the kinetics of fusion activity data (see **Methods**). **g,** Initial velocity of the S-mediated fusion. In **b,c**, statistically significant differences versus the D614G mutant-infected culture (*, *P* < 0.05) and uninfected culture (#, *P* < 0.05) were determined by the Mann-Whitney U test. In **d**, a statistically significant difference (*, *P* < 0.05) versus the D614G S was determined by Student’s *t* test. In **g,** statistically significant differences (*, *P* < 0.05) were determined by two-sided Welch’s *t* test.

To clearly observe the syncytia formation, we further generated the GFP-expressing replication-competent D614G and D614G/P681R viruses. The levels of viral RNA in the supernatant and GFP-positive cells were similar in Vero, VeroE6/TMPRSS2 and Calu-3 cells (Extended Data Fig. 4). However, at 24 hpi, significantly larger GFP-positive adherent syncytia were observed in the VeroE6/TMPRSS2 cells infected with the GFP-expressing D614G/P681R virus (Fig. 3c). Additionally, the size of GFP-positive floating syncytia at 72 hpi in the VeroE6/TMPRSS2 cells infected with GFP-expressing D614G/P681R virus was significantly bigger than that with GFP-expressing D614G virus (Extended Data Fig. 5). Moreover, GFP-positive syncytia were observed in the D614G/P681R-infected but not in the D614G-infected Calu-3 cells at 72 hpi (Extended Data Fig. 4c). These results suggest that the feature of the B.1.617.2/Delta virus observed in *in vitro* cell culture experiments, particularly forming larger syncytia (Fig. 2b and Extended Data Fig. 2), is well reproduced by the insertion of P681R mutation.

To directly investigate the effect of P681R mutation on the cleavage of SARS-CoV-2 S protein, we prepared the HIV-1-based pseudoviruses carrying the P681R mutation. Western blotting of the pseudoviruses prepared showed that the level of cleaved S2 subunit was significantly increased by the P681R mutation (Extended Data Fig. 6a), suggesting that the P681R mutation facilitates the furin-mediated cleavage of SARS-CoV-2 S protein. We then performed the single-round pseudovirus infection assay using the target HOS cells with or without TMPRSS2 expression. The infectivity of both the D614G and the D614G/P681R pseudoviruses increased approximately 10-fold by the expression of TMPRSS2 in target cells (Extended Data Fig. 6b). However, the infectivity of the D614G and the D614G/P681R pseudoviruses were comparable regardless of TMPRSS2 expression (Extended Data Fig. 6b). These data suggest that the P681R mutation does not affect the infectivity of viral particles.

We next addressed the effect of P681R mutation on viral fusogenicity by cell-based fusion assay. In the effector cells (i.e., S-expressing cells), although the expression level of the D614G/P681R S protein was comparable to that of the D614G S, the level of the cleaved S2 subunit of the D614G/P681R mutant was significantly higher than that of the D614G S (Fig. 3d). Consistent with the results in the pseudovirus assay (Extended Data Fig. 6), these results suggest that P681R mutation facilitates the S cleavage. Flow cytometry showed that the surface expression level of the D614G/P681R S was significantly lower than the D614G S (Extended Data Fig. 7). Nevertheless, the cell-based fusion assay using the target cells without TMPRSS2 demonstrated that the D614G/P681R S is 2.1-fold more fusogenic than the D614G S with a statistical significance (*P* = 0.0002 by Welch’s t test) (Fig. 3e). Moreover, a mathematical modeling analysis of the fusion assay data showed that the initial fusion velocity of the D614G/P681R S (0.83 ± 0.03 per hour) is significantly (2.8-fold) faster than that of the D614G S (0.30 ± 0.03 per hour; *P* = 4.0 × 10^-6^ by Welch’s t test) (Fig. 3f, g). These data suggest that the P681R mutation enhances and accelerates the SARS-CoV-2 S-mediated fusion. Furthermore, when we use the target cells with TMPRSS2 expression, both the fusion efficacy (∼1.2-fold) and initial fusion velocity (∼2.0-fold) were increased in both the D614G and D614G/P681R S proteins (Fig. 3f, g). These results suggest that TMPRSS2 facilitates the fusion mediated by SARS-CoV-2 S and human ACE2, while the TMPRSS2-dependent acceleration and promotion of viral fusion is not specific for the P681R mutant.

### Resistance to NAb-mediated antiviral immunity by the P681R mutation

The resistance to the NAb in the sera of COVID-19 convalescents and vaccinated individuals is a hallmark characteristic of the VOCs (reviewed in ^20, 21^), and it has been recently showed that the B.1.617.2/Delta variant is relatively resistant to the vaccine-induced neutralisation^17, 18^. To ask whether the P681R mutation contributes to this virological phenotype, we performed the neutralisation assay. The D614G/P681R pseudovirus was partially (1.2-1.5-fold) resistant to the three monoclonal antibodies targeting the receptor binding domain of SARS-CoV-2 S protein (Extended Data Fig. 8a). Additionally, the neutralisation experiments using the 19 sera of second BNT162b2 vaccination showed that the D614G/P681R pseudovirus is significantly resistant to the vaccine-induced NAbs compared to the D614G pseudovirus (*P* < 0.0001 by Wilcoxon matched-pairs signed rank test; Extended Data Fig. 8b and 9). These results suggest that the P681R-bearing pseudovirus is relatively resistant to NAbs. Notably, in contrast to the neutralising activity against cell-free viruses, the SARS-CoV-2 S-based fusion assay showed that cell-cell infection is strongly resistant to the NAbs and the insensitivity to the NAbs on cell-cell infection is not dependent on the P681R mutation (Extended Data Fig. 8c). Altogether, these findings suggest that the P681R mutation confers the NAbs resistance upon cell-free viral particles and cell-cell infection is resistant to the NAb-mediated antiviral action compared to cell-free infection.

### P681R mutation as the determinant of higher pathogenicity of the Delta variant

To assess the impact of the P681R mutation on viral replication and the pathogenicity of SARS-CoV-2, Syrian hamsters were intranasally infected with the D614G and D614G/P681R viruses. The D614G-infected hamsters exhibited no weight loss, although a slight decrease in body weight was observed for one of the animals by 7 dpi (5.0%) (Fig. 4a). In contrast, all of hamsters infected with the D614G/P681R virus experienced a gradual body weight loss and the animals showed a significant weight loss of 4.7-6.9% at 7 dpi compared to the D614G virus (*P* = 0.011, Fig. 4a). We also assessed pulmonary function in infected hamsters by measuring enhanced pause (PenH), which is a surrogate marker for bronchoconstriction or airway obstruction, by using a whole-body plethysmography system. Syrian hamsters infected with the D614G and D614G/P681R viruses showed the increases in the lung PenH value (Fig. 4b). At 7 dpi, the D614G/P681R-infected animals had significantly higher PenH values compared with those of the D614G-infected animals (*P* = 0.043). At 3 dpi, both viruses replicated efficiently in the lungs and nasal turbinates of the infected animals and no significant difference in viral replication was observed between the two groups (Fig. 4c). At 7 dpi, no differences in viral titres in the nasal turbinates were found between the two groups; however, the lung titres in the D614G/P681R-infected group were significantly higher than those in the D614G-infected groups (*P* = 0.0013, Fig. 4c).

**Fig. 4.**
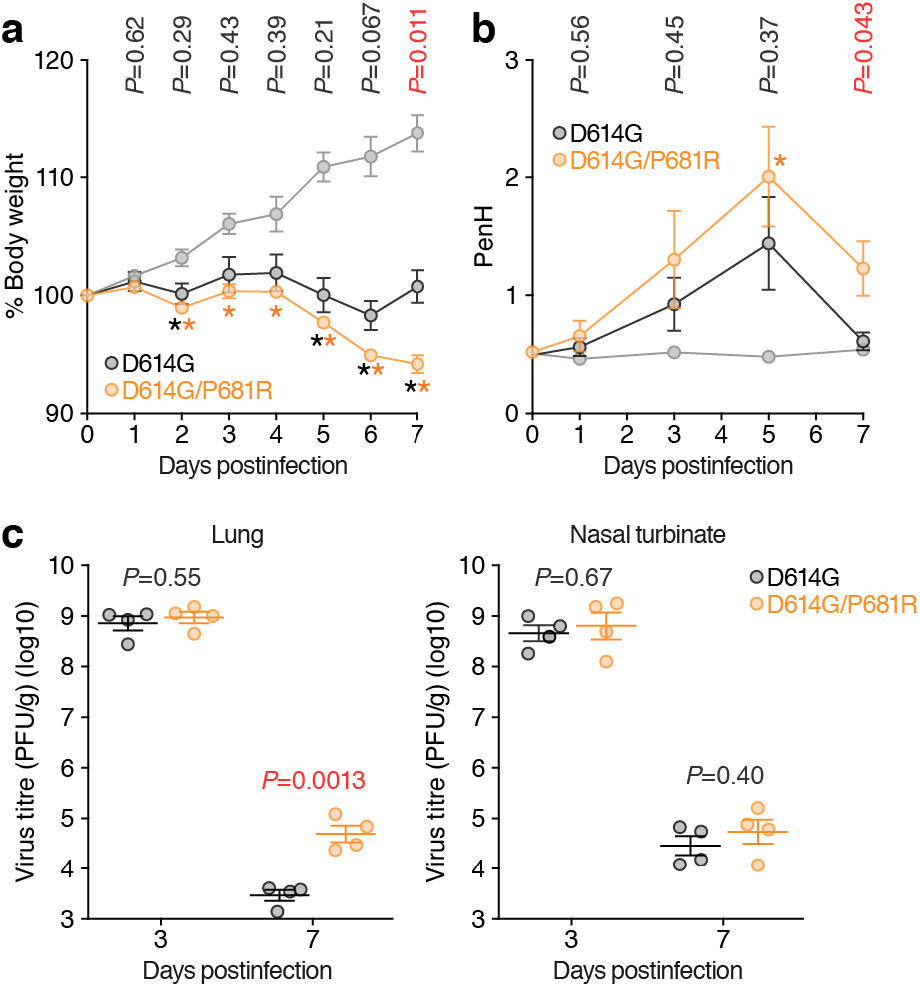
Enhanced pathogenicity by the P681R mutation in hamsters. Syrian hamsters were intranasally inoculated with 10^4^ TCID_50_ (in 30 μl) of the D614G and D614G/P681R viruses. **a**, Body weight changes in hamsters after viral infection. Body weights of virus-infected (n = 4 each) and uninfected hamsters (n = 3) were monitored daily for 7 days. **b**, Pulmonary function analysis in infected hamsters. Enhanced pause (PenH), which is a surrogate marker for bronchoconstriction or airway obstruction, was measured by using whole-body plethysmography. **c**, Virus replication in infected hamsters. Four hamsters per group were euthanized at 3 and 7 dpi for virus titration. Virus titres in the lungs (left) nasal turbinates (right) were determined by the plaque assay using VeroE6/TMPRSS2 cells. Points indicate data from individual Syrian hamsters. Statistically significant differences were determined by the Mann-Whitney U test, and those versus uninfected hamsters (*, *P* < 0.05) are indicated by asterisks. The *P* value between the D614G and the D614G/P681R at each dpi is indicated in the figure.

## Discussion

Previous studies have demonstrated the close association of the FCS in the SARS-CoV-2 S protein with viral replication mode and it is dependent on TMPRSS2. Johnson et al. and Peacock et al. showed that the loss of FCS results in the increase of viral replication efficacy in Vero cells while the attenuation of viral growth in the Vero cells expressing TMPRSS2^22, 23^. On the contrary, here we showed that the replication efficacy of the B.1.617.2/Delta variant was severely decreased in Vero cells compared to VeroE6/TMPRSS2 cells. More importantly, although the FCS-deleted SARS-CoV-2 is less pathogenic than the parental virus^23^, we revealed that the B.1.617.2/Delta variant as well as the P681R-harbouring virus exhibit higher pathogenicity. These findings suggest that the enhanced viral fusogenicity, which is triggered by the P681R mutation, closely associates with viral pathogenicity.

It is evident that most VOCs considered so far have acquired mutations in their S proteins, particularly in the RBD and N-terminal domain, to evade NAbs^20, 21, 24^. In sharp contrast, here we demonstrated that the B.1.617.2/Delta variant has acquired a unique strategy to facilitate infection and evade antiviral immunity. The P681R mutation that is highly conserved in this lineage enhances the efficacy of viral fusion and further accelerates its speed of action. The P681R-mediated rapid kinetics of viral fusion may attribute to not only immune evasion but also possibly feasible the infection to exposed individuals.

Consistent with previous reports^25, 26^, here we showed that the cell-cell infection mediated by the SARS-CoV-2 S protein is resistant to NAbs. The effect of NAbs against cell-cell infection has been well studied in HIV-1 (*Retroviridae*) infection, and it is well known that cell-cell infection is relatively more resistant to NAbs compared to cell-free infection (reviewed in ^27–29^). The resistance of cell-cell spread against NAbs is not limited to HIV-1 but has been observed in the other viruses such as vaccinia virus (*Poxviridae*)^30^ and hepatitis C virus (*Flaviviridae*)^31^, suggesting that cell-cell infection is a common strategy for a variety of viruses to evade antiviral humoral immunity. The fact that the B.1.617.2/Delta variant as well as the P681R mutant efficiently form syncytia and the P681R mutant accelerates and promotes cell-cell fusion suggests that switching the preference of viral replication mode from cell-free infection to cell-cell infection may be a unique strategy of the B.1.617.2/Delta variant to evade antiviral immunity.

Although the P681R mutant is highly fusogenic, the virus harbouring the P681R mutation did not necessarily show higher growth compared to the parental virus in *in vitro* cell cultures. Regarding this, the HIV-1 variants with higher fusogenicity have been isolated from AIDS patients, but the enhanced fusogenicity does not promote viral replication in *in vitro* cell cultures^32^. Similarly, the measles virus (*Paramyxoviridae*) harbouring the deficient mutation in viral matrix protein^33^ and substitution mutations in viral fusion protein^34, 35^ are highly fusogenic and efficiently expands via cell-cell fusion. However, the growth kinetics of these mutated measles virus with higher fusogenicity in *in vitro* cell cultures is less efficient than the parental virus^33^. Therefore, the discrepancy between the efficacy of viral growth in *in vitro* cell cultures and viral fusogenicity is not specific for SARS-CoV-2. Rather, the higher fusogenicity is associated with the severity of viral pathogenicity such as HIV-1 encephalitis^36^ and fatal subacute sclerosing panencephalitis, which is caused by measles virus infection in brain^34, 35^. Consistently, here we showed that the B.1.617.2/Delta variant as well as the P681R mutant exhibited a higher fusogenicity *in vitro* and enhanced pathogenicity *in vivo*. Our data suggest that the higher COVID-19 severity and unusual symptoms caused by the B.1.617.2/Delta variant^15, 16^ are partly due to the higher fusogenicity caused by the P681R mutation. Switching viral infection mode by the P681R mutation may relate to the severity and/or unusual outcome of viral infection, therefore, the epidemic of the SARS-CoV-2 variants harbouring the P681R mutation should be surveyed in depth.

## Author Contributions

A.S., T.Irie, R.Suzuki, H.N., K.U., Y.Kosugi, E.P.B., Y.L.T., R.Shimizu, K.Shimizu, R.K., T.Ikeda, T.F. and K.Sato performed the experiments.

R.Suzuki, T.M., K.I.-H., M.Ito, S.Y., M.Imai, K.Yoshimatsu, T.F. and Y.Kawaoka performed animal experiments.

K.Sadamasu, S.O., T.S., K.T., I.Y., H.A., M.N., and K.Yoshimura prepared experimental materials.

J.W. and S.N. performed molecular phylogenetic analysis.

K.Shirakawa, Y.Kazuma, R.N., Y.H., and A.T.-K. collected clinical samples.

A.S., T.Irie, M.Imai, S.N., T.Ikeda, T.F. Y.Kawaoka and K.S. designed the experiments and interpreted the results.

K.Sato wrote the original manuscript.

All authors reviewed and proofread the manuscript.

The Genotype to Phenotype Japan (G2P-Japan) Consortium contributed to the project administration.

## Consortia

The Genotype to Phenotype Japan (G2P-Japan) Consortium: Mika Chiba, Hirotake Furihata, Haruyo Hasebe, Kazuko Kitazato, Haruko Kubo, Naoko Misawa, Nanami Morizako, Akiko Oide, Mai Suganami, Miyoko Takahashi, Kana Tsushima, Miyabishara Yokoyama, Yue Yuan

## Acknowledgments

We would like to thank all members belonging to The Genotype to Phenotype Japan (G2P-Japan) Consortium. We thank Dr. Jin Gohda (The University of Tokyo, Japan) for providing reagents. An anti-HIV-1 p24 Monoclonal antibody (clone 183-H12-5C, Cat# ARP-3537) was obtained through the NIH HIV Reagent Program, NIAID, NIH (contributed by Drs. Bruce Chesebro and Kathy Wehrly). The super-computing resource was provided by Human Genome Center at The University of Tokyo and the NIG supercomputer at ROIS National Institute of Genetics.

This study was supported in part by AMED Research Program on Emerging and Re-emerging Infectious Diseases 20fk0108163 (to A.S.), 20fk0108401 (to T.F.), 21fk0108617 (to T.F.), 19fk0108113 (to Y.Kawaoka), JP20fk0108412 (to Y.Kawaoka), 20fk0108146 (to K.Sato), 20fk0108270 (to K.Sato) and 20fk0108413 (to T.Ikeda, S.N. and K.Sato); AMED Research Program on HIV/AIDS 21fk0410033 (to A.S.) and 21fk0410039 (to K.Sato); AMED Japan Program for Infectious Diseases Research and Infrastructure 20wm0325009 (to A.S.), 21wm0325009 (to A.S.) and 21wm0125002 (to Y.Kawaoka); JST A-STEP JPMJTM20SL (to T.Ikeda); JST SICORP (e-ASIA) JPMJSC20U1 (to K.Sato); JST SICORP JPMJSC21U5 (to K.Sato), JST CREST JPMJCR20H6 (to S.N.) and JPMJCR20H4 (to K.Sato); JSPS KAKENHI Grant-in-Aid for Scientific Research C 19K06382 (to A.S.), 18K07156 (to K.T.) and 21K07060 (to K.T.), Scientific Research B 18H02662 (to K.Sato) and 21H02737 (to K.Sato); JSPS Fund for the Promotion of Joint International Research (Fostering Joint International Research) 18KK0447 (to K.Sato); JSPS Core-to-Core Program JPJSCCA20190008 (A. Advanced Research Networks) (to K.Sato); JSPS Research Fellow DC1 19J20488 (to I.K.); JSPS Leading Initiative for Excellent Young Researchers (LEADER) (to T.Ikeda); ONO Medical Research Foundation (to K.Sato); Ichiro Kanehara Foundation (to K.Sato); Lotte Foundation (to K.Sato); Mochida Memorial Foundation for Medical and Pharmaceutical Research (to K.Sato); Daiichi Sankyo Foundation of Life Science (to K.Sato); Sumitomo Foundation (to K.Sato); Uehara Foundation (to K.Sato); Takeda Science Foundation (to T.Ikeda and K.Sato); The Tokyo Biochemical Research Foundation (to K.Sato); Mitsubishi Foundation (to T.Ikeda); Shin-Nihon Foundation of Advanced Medical Research (to T.Ikeda); Tsuchiya Foundation (to T.Irie); a Grant for Joint Research Projects of the Research Institute for Microbial Diseases, Osaka University (to A.S.); an intramural grant from Kumamoto University COVID-19 Research Projects (AMABIE) (to T.Ikeda); Intercontinental Research and Educational Platform Aiming for Eradication of HIV/AIDS (to T.Ikeda); and Joint Usage/Research Center program of Institute for Frontier Life and Medical Sciences, Kyoto University (to K.Sato).

---

**Extended Data Fig. 1.**
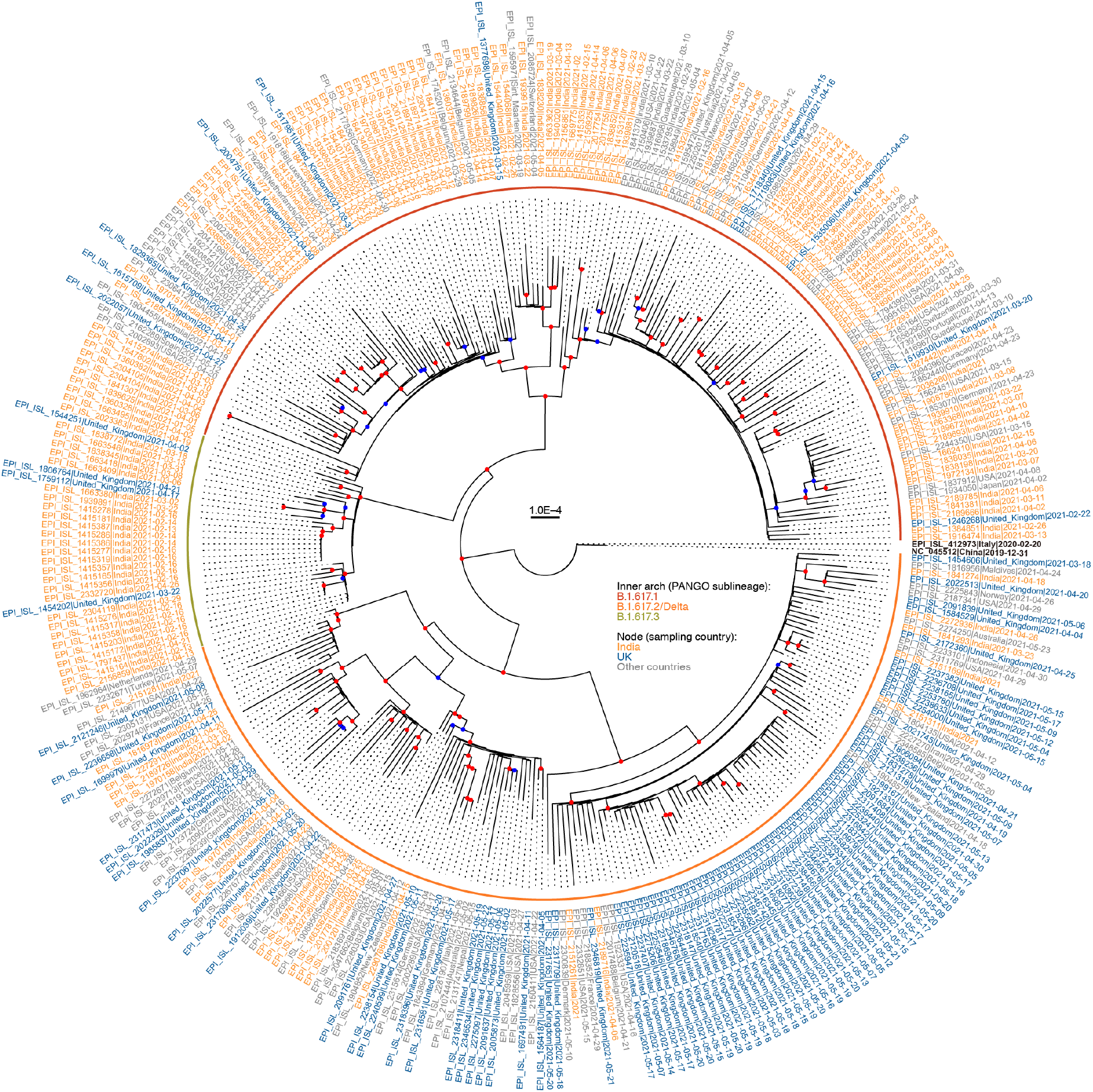
A maximum-likelihood based phylogenetic tree of the representative 334 SARS-CoV-2 sequences that belong to the B.1.617 lineage. GISAID ID, exposure country, and sampling date were noted in each terminal node. The country isolated (India, UK, or the other countries) and the PANGO sublineage are labeled by colors as indicated in the figure. Red or blue circle on the branch was shown in each internal node if the bootstrap value was ≥ 80 or ≥ 50 (n = 1,000).

**Extended Data Fig. 2.**
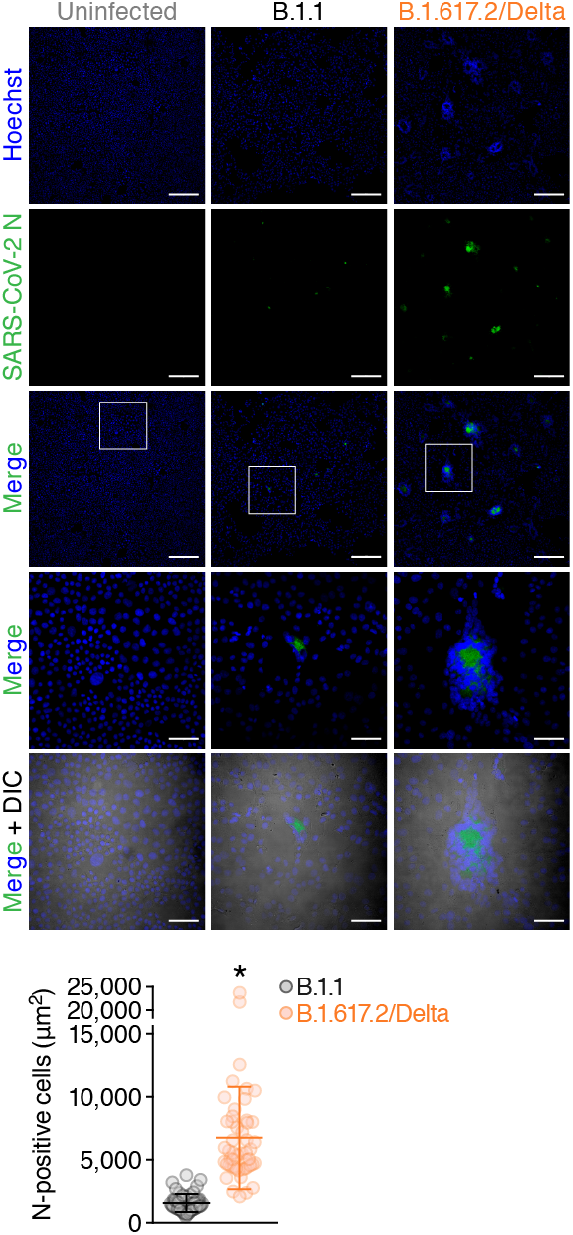
Immunofluorescence staining of B.1.617.2/Delta-infected VeroE6/TMPRSS2 cells. VeroE6/TMPRSS2 cells infected with the viruses indicated [multiplicity of infection (MOI) 0.01] were stained with anti-SARS-CoV-2 nucleocapsid (N) (green) and Hoechst (blue). (Top) Representative images at 48 hpi are shown. Areas enclosed with squares are enlarged in the bottom panels. DIC, differential interference contrast. Bars, 200 μm for low magnification panels; 50 μm for high magnification panels. (Bottom) The area of N-positive cells in B.1.1-infected (n = 50) and B.1.617.2/Delta-infected (n = 50) cultures are shown. A statistically significant difference versus the B.1.1-infected culture (*, *P* < 0.05) was determined by the Mann-Whitney U test.

**Extended Data Fig. 3.**
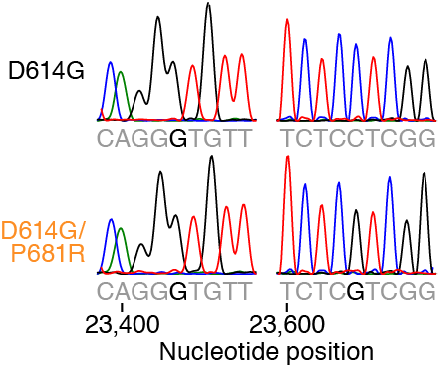
Chromatograms of the mutated regions of SARS-CoV-2 viruses artificially generated by reverse genetics. Chromatograms of nucleotide positions 23,399-23,407 (left) and 23,600-23,608 (right) of parental SARS-CoV-2 (strain WK-521, PANGO lineage A; GISAID ID: EPI_ISL_408667) and the D614G (A23403G in nucleotide) and P681R (C23604G in nucleotide) mutation are shown.

**Extended Data Fig. 4.**
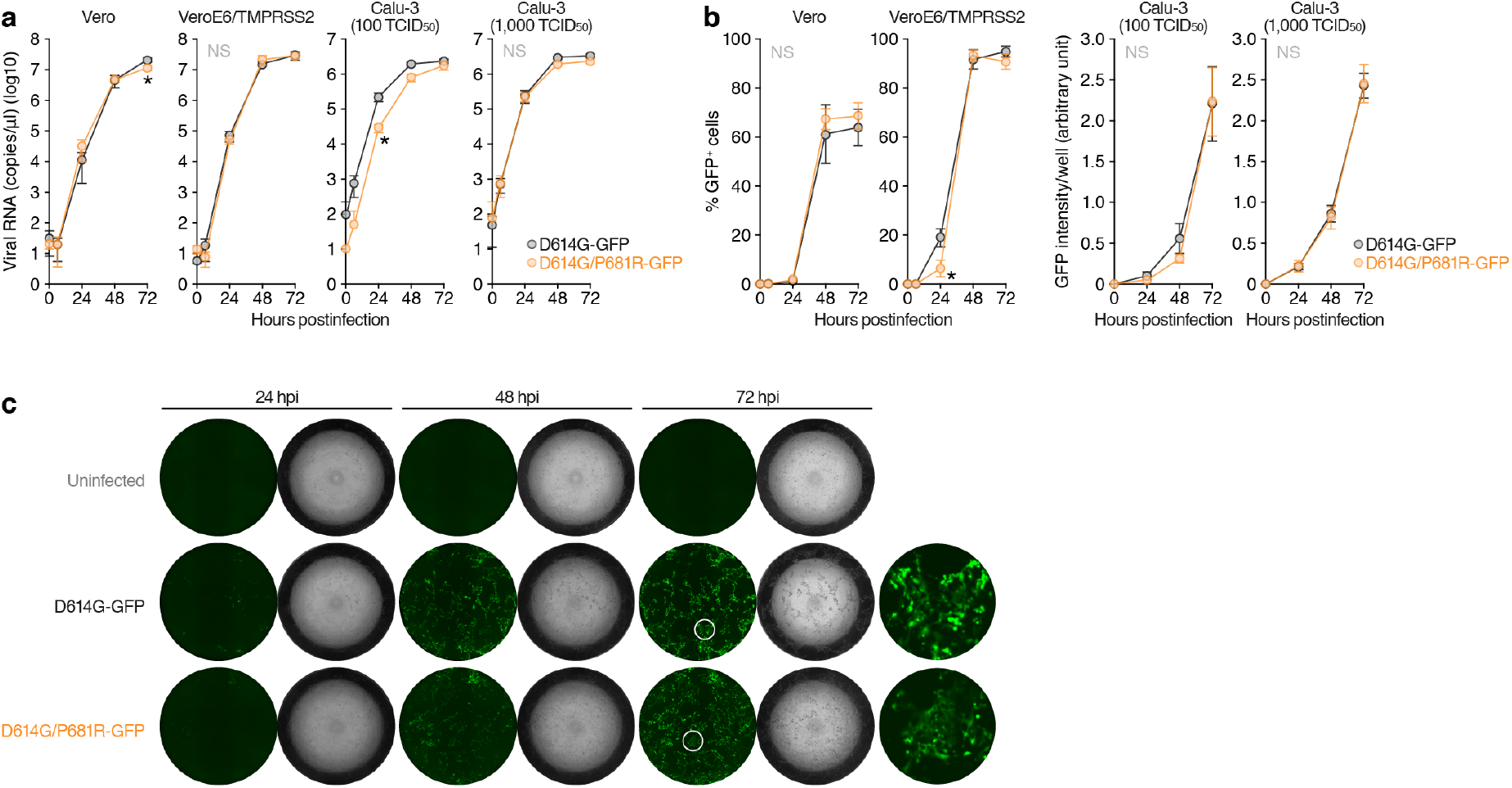
Growth kinetics of artificially generated GFP-expressing viruses. The GFP-expressing D614G and D614G/P681R mutant viruses were generated by reverse genetics. These viruses (100 TCID_50_ for Vero and VeroE6/TMPRSS2 cells, 100 or 1,000 TCID_50_ for Calu-3 cells) were inoculated. The copy number of viral RNA in the culture supernatant (**a**) and the level of GFP-positive cells (the percentage of GFP-positive cells, for Vero and VeroE6/TMPRSS2 cells; the GFP intensity per well, for Calu-3 cells) (**b**) are shown. (**c**) Representative images of the Calu-3 cells infected with GFP-expressing viruses (100 TCID_50_). Areas enclosed with circles are enlarged in the right panels. Assays were performed in quadruplicate. Statistically significant differences (*, *P* < 0.05) versus the D614G virus were determined by Student’s *t* test. NS, no statistical significance.

**Extended Data Fig. 5.**
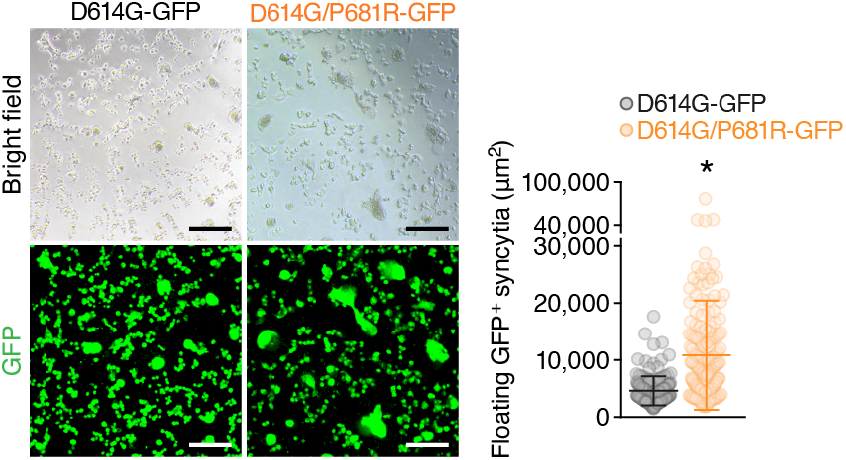
Syncytia formation in VeroE6/TMPRSS2 cells infected with GFP-expressing viruses. (Left) Floating syncytia in VeroE6/TMPRSS2 cells infected with the GFP-expressing D614G and D614G/P681R mutant viruses (100 TCID_50_) at 72 hpi are shown. Bars, 100 μm. (Right) The size of adherent GFP^+^ syncytia in the D614G mutant-infected (n = 147) and the D614G/P681R mutant-infected (n = 171) cultures are shown. A statistically significant difference versus the D614G mutant-infected culture (*, *P* < 0.05) was determined by the Mann-Whitney U test.

**Extended Data Fig. 6.**
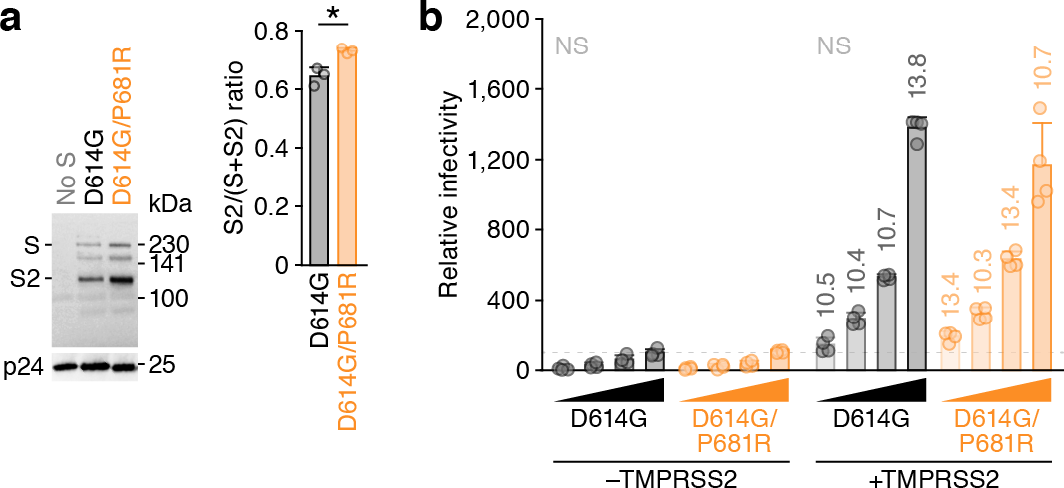
Infectivity of the P681R-bearing pseudovirus. **a,** Western blotting of pseudoviruses. (Left) Representative blots of SARS-CoV-2 full-length S and cleaved S2 proteins as well as HIV-1 p24 capsid as an internal control. kDa, kilodalton. (Right) The ratio of S2 to the full-length S plus S2 proteins on pseudovirus particles. **b,** Pseudovirus assay. The HIV-1-based reporter virus pseudotyped with the SARS-CoV-2 S D614G or D614G/P681R was inoculated into HOS-ACE2 cells or HOS-ACE2/TMPRSS2 cells at 4 different doses (125, 250, 500 and 1,000 ng HIV-1 p24 antigen). Percentages of infectivity compared to the virus pseudotyped with parental S D614G (1,000 ng HIV-1 p24) in HOS-ACE2 cells are shown. The numbers on the bars of the HOS-ACE2/TMPRSS2 cell data indicate the fold change versus the HOS-ACE2 cell data. Assays were performed in quadruplicate. NS, no statistical significance.

**Extended Data Fig. 7.**
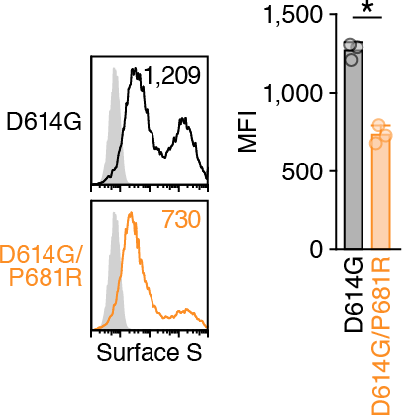
Flow cytometry of the S-expressing cells. (Left) Representative histogram of the S protein expression on the cell surface. The number in the histogram indicates the mean fluorescence intensity (MFI). (Right) The MFI of surface S on the S-expressing cells. A statistically significant difference (*, *P* < 0.05) versus the D614G S was determined by Student’s *t* test.

**Extended Data Fig. 8.**
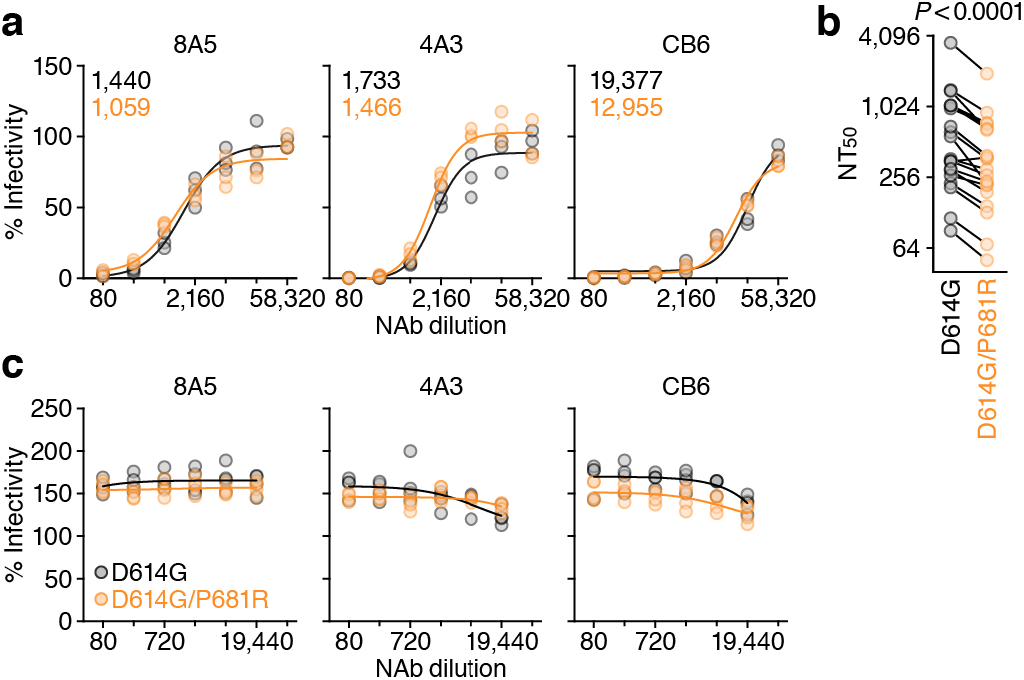
Association of the P681R mutation on the sensitivity to NAbs. Neutralisation assay was performed by using three RBD-targeting monoclonal antibodies (clones 8A5, 4A3 and CB6) (**a** and **c**) and 19 vaccinated sera (**b**). NAbs were used for the pseudovirus assay (**a** and **b**) and the S-based fusion assay (**c**). Pseudoviruses and effector cells (S-expressing cells) were treated with serially diluted NAbs or sera as described in **Methods**. The raw data of panel **b** is shown in Extended Data Fig. 9. NT_50_, 50% neutralisation titre. In **a**, the NT_50_ values of the D614G S (black) and D614G/P681R S (orange) are indicated. In **b**, a statistically significant difference versus the D614G virus was determined by Wilcoxon matched-pairs signed rank test.

**Extended Data Fig. 9.**
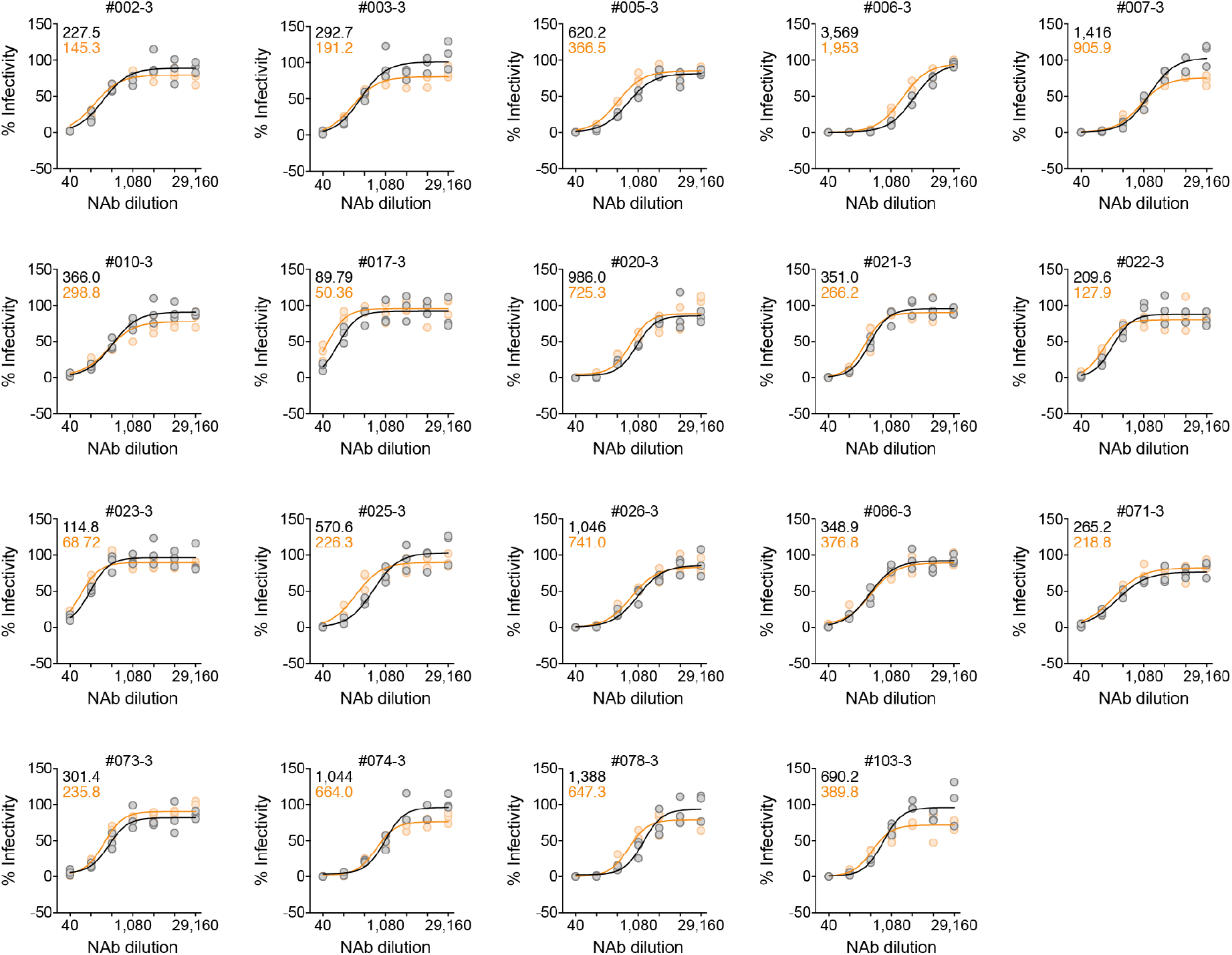
Neutralisation assay using 19 vaccinated sera. Nineteen vaccinated sera were used for the neutralisation assay. The NT_50_ values of respective serum against the D614G S (black) and D614G/P681R S (orange) are indicated in each panel. The NT_50_ values are summarized in Extended Data Fig. 8b.

**Extended Data Table 1.** Number of daily deposited sequences in GISAID.

**Extended Data Table 2.** Percentage of the mutations detected in the S protein of the B.1.617 lineage.

**Extended Data Table 3.** The SARS-CoV-2 genomic region encoded by each template and the primers used for the preparation of each fragment for CPER.

## Methods

### Ethics Statement

For virus isolation, this study was approved by the Institutional Review Board of Tokyo Metropolitan Institute of Public Health, according to the Declaration of Helsinki 2013 (approval number 3KenKenKen-466). For the use of human specimen, all protocols involving human subjects recruited at Kyoto University were reviewed and approved by the Institutional Review Boards of Kyoto University (approval number G0697). All human subjects provided written informed consent. All experiments with hamsters were performed in accordance with the Science Council of Japan’s Guidelines for Proper Conduct of Animal Experiments. The protocols were approved by the Institutional Animal Care and Use Committee of National University Corporation Hokkaido University (approval number 20-0123) and the Animal Experiment Committee of the Institute of Medical Science, the University of Tokyo (approval number PA19-75).

### Collection of BNT162b2-Vaccinated Sera

Peripheral blood were collected four weeks after the second vaccination of BNT162b2 (Pfizer-BioNTech), and the sera of 19 vaccinees (average age: 38, range: 28-59, 26% male) were isolated from peripheral blood. Sera were inactivated at 56°C for 30 min and stored at –80°C until use.

### Cell Culture

HEK293 cells (a human embryonic kidney cell line; ATCC CRL-1573), HEK293T cells (a human embryonic kidney cell line; ATCC CRL-3216), and HOS cells (a human osteosarcoma cell line; ATCC CRL-1543) were maintained in Dulbecco’s modified Eagle’s medium (high glucose) (Wako, Cat# 044-29765) containing 10% fetal bovine serum (FBS) and 1% PS. Vero cells [an African green monkey (*Chlorocebus sabaeus*) kidney cell line; JCRB0111] were maintained in Eagle’s minimum essential medium (Wako, Cat# 051-07615) containing 10% FBS and 1% PS. VeroE6/TMPRSS2 cells [an African green monkey (*Chlorocebus sabaeus*) kidney cell line; JCRB1819]^37^ were maintained in Dulbecco’s modified Eagle’s medium (low glucose) (Wako, Cat# 041-29775) containing 10% FBS, G418 (1 mg/ml; Nacalai Tesque, Cat# G8168-10ML) and 1% PS. Calu-3 cells (a human lung epithelial cell line; ATCC HTB-55) were maintained in Minimum essential medium Eagle (Sigma-Aldrich, cat# M4655-500ML) containing 10% FCS and 1% PS. HOS-ACE2/TMPRSS2 cells, the HOS cells stably expressing human ACE2, was prepared as described previously^8, 38^. HEK293-C34 cells, the *IFNAR1* KO HEK293 cells expressing human ACE2 and TMPRSS2 by doxycycline treatment^39^, were maintained in Dulbecco’s modified Eagle’s medium (high glucose) (Sigma-Aldrich, Cat# R8758-500ML) containing 10% FBS, 10 μg/ml blasticidin (InvivoGen, Cat# ant-bl-1) and 1% PS.

### Animal experiments

Syrian hamsters (Male, 4 weeks old) were purchased from Japan SLC Inc. (Shizuoka, Japan). Baseline body weights were measured before infection. For the virus infection in Fig. 2c**, d**, hamsters were euthanised by intramuscular injection of a mixture of 0.15 mg/kg medetomidine hydrochloride (Domitor^®^, Nippon Zenyaku Kogyo), 2.0 mg/kg midazolam (Dormicum^®^, Maruishi Pharmaceutical) and 2.5 mg/kg butorphanol (Vetorphale^®^, Meiji Seika Pharma). The B.1.1 or B.1.167.2/Delta viruses (10^5^ TCID_50_ in 100 µl) was intranasally infected under anesthesia. Body weights were measured and oral swabs were collected under anesthesia with isoflurane (Sumitomo Dainippon Pharma) daily. For the virus infection in Fig. 4, four hamsters per group were intranasally inoculated with the D614G or the D614G/P681R viruses (10^4^ TCID_50_ in 30 μl) under isoflurane anesthesia. Body weight was monitored daily for 7 days. For virological examinations, four hamsters per group were intranasally infected with the D614G or the D614G/P681R viruses (10^4^ TCID_50_ in 30 μl); at 3 and 7 dpi, the animals were euthanized and nasal turbinates and lungs were collected. The virus titers in the nasal turbinates and lungs were determined by use of plaque assays on VeroE6/TMPRSS2 cells.

### Lung function

Respiratory parameters were measured by using a whole-body plethysmography system (PrimeBioscience) according to the manufacturer’s instructions. In brief, hamsters were placed in the unrestrained plethysmography chambers and allowed to acclimatize for 1 m before data were acquired over a 3-min period by using FinePointe software.

### Viral Genomes

All SARS-CoV-2 genome sequences and annotation information used in this study were downloaded from GISAID (https://www.gisaid.org) as of May 31, 2021 (1,761,037 sequences). We first excluded the genomes with non-human hosts. We obtained SARS-CoV-2 variants belonging to the B.1.617 lineage based on the PANGO annotation (i.e. sublineages B.1.617.1, B.1.617.2/Delta, or B.1.617.3) for each sequence in the GISAID metadata. Note that only one variant belonging to the B.1.617 lineage (GISAID ID: EPI_ISL_1544002 isolated in India on February 25, 2021) was not used in the analysis because the variant is not assigned any three sublineages possibly due to 212 undetermined nucleotides in the genome. To infer epidemiology of the B.1.617 lineage (Fig. 1b-1f), we excluded genomes that sampling date information are not available, and collected 2,855, 13,821, or 83 sequences belonging to the B.1.617.1, B.1.617.2/Delta, or B.1.617.3 sublineage, respectively.

### Phylogenetic Analyses

To infer the phylogeny of the B.1.617 sublineages, we screened SARS-CoV-2 genomes by removing genomes containing undetermined nucleotides at coding regions. Since the number of genomes belonging to the sublineage B.1.617.1 or B.1.617.2/Delta are large (i.e. 894 or 6152 sequences, respectively), we used 150 sequences randomly chosen for each sublineage. For the B.1.617.3 sublineage, 32 genomes were used. We used Wuhan-Hu-1 strain isolated in China on December 31, 2019 (GenBank ID: NC_045512.2 and GISAID ID: EPI_ISL_402125) and LOM-ASST-CDG1 strain isolated Italy on February 20, 2020 (GISAID ID: EPI_ISL_412973) as an outgroup. We then collected 334 representative SARS-CoV-2 sequences, and aligned entire genome sequences by using the FFT-NS-1 program in MAFFT suite v7.407^40^. All sites with gaps in alignment are removed, and the total length of alignment is 29,085 nucleotides. Maximum likelihood tree was generated by IQ-TREE 2 v2.1.3 software with 1,000 bootstraps^41^. GTR+G substitution model is utilized based on BIC criterion.

### SARS-CoV-2 Preparation and Titration

A B.1.617.2/Delta isolate (GISAID ID: EPI_ISL_2378732) and a D614G-bearing B.1.1 isolate (GISAID ID: EPI_ISL_479681) were isolated from SARS-CoV-2-positive individuals in Japan. Briefly, 100 μl of the nasopharyngeal swab obtained from SARS-CoV-2-positive individuals were inoculated into VeroE6/TMPRSS2 cells in the biosafety level 3 laboratory. After the incubation at 37°C for 15 minutes, a maintenance medium supplemented with Eagle’s minimum essential medium (FUJIFILM Wako Pure Chemical Corporation, Cat# 056-08385) including 2% FBS and 1% PS was added, and the cells were cultured at 37°C under 5% CO_2_. The cytopathic effect (CPE) was confirmed under an inverted microscope (Nikon), and the viral load of the culture supernatant in which CPE was observed was confirmed by real-time RT-PCR. To determine viral genome sequences, RNA was extracted from the culture supernatant using QIAamp viral RNA mini kit (Qiagen, Qiagen, Cat# 52906). cDNA library was prepared by using NEB Next Ultra RNA Library Prep Kit for Illumina (New England Biolab, Cat# E7530) and whole genome sequencing was performed by Miseq (Illumina).

To prepare the working virus, 100 μl of the seed virus was inoculated into VeroE6/TMPRSS2 cells (5,000,000 cells in a T-75 flask). At one hour after infection, the culture medium was replaced with Dulbecco’s modified Eagle’s medium (low glucose) (Wako, Cat# 041-29775) containing 2% FBS and 1% PS; at 2-3 days postinfection, the culture medium was harvested and centrifuged, and the supernatants were collected as the working virus.

The titre of the prepared working virus was measured as 50% tissue culture infectious dose (TCID_50_). Briefly, one day prior to infection, VeroE6/TMPRSS2 cells (10,000 cells/well) were seeded into a 96-well plate. Serially diluted virus stocks were inoculated to the cells and incubated at 37°C for 3 days. The cells were observed under microscopy to judge the CPE appearance. The value of TCID_50_/ml was calculated with the Reed–Muench method^42^.

### SARS-CoV-2 Infection

One day prior to infection, Vero cells (10,000 cells), VeroE6/TMPRSS2 cells (10,000 cells) and Calu-3 cells (10,000 cells) were seeded into a 96-well plate. SARS-CoV-2 was inoculated and incubated at 37°C for 1 h. The infected cells were washed, and 180 µl of culture medium was added. The culture supernatant (10 µl) was harvested at indicated time points and used for real-time RT-PCR to quantify the viral RNA copy number. To monitor the syncytia formed in infected cell culture, the bright-field photos were obtained using ECLIPSE Ts2 (Nikon). The size of floating syncytia was measured by “quick selection tool” in Photoshop CS5 (Adobe) as pixel, and the area of floating syncytia was calculated from the pixel value. As for the GFP-expressing recombinant viruses (Extended Data Fig. 4b, c), the bright-field and green fluorescence photos were obtained using an All-in-One Fluorescence microscope BZ-X800 (Keyence) at the indicated time points, and the GFP-fluorescence intensity was analyzed by a BZ-X800 Analyzer (Keyence).

### Immunofluorescence Staining

One day prior to infection, VeroE6/TMPRSS2 cells (200,000 cells) were seeded on the coverslips put in 12-well plate and were infected with SARS-CoV-2 (2,000 TCID_50_). At 48 hours postinfection, the cells were fixed with 4% paraformaldehyde in phosphate buffer saline (PBS) (Nacalai Tesque, Cat# 09154-85) for 10 min at room temperature. The fixed cells were permeabilized with 0.1% Triton X-100 in PBS for 10 min, blocked with 10% FBS in PBS for overnight at 4°C, and then stained using mouse anti-SARS-CoV-2 N monoclonal antibody (GeneTex, Cat# GTX632269) for 1 h. After washing three times with PBS, cells were incubated with an Alexa 488-conjugated anti-mouse IgG antibody (Jackson ImmunoResearch, Cat# 015-540-003) for 1 h. Nuclei were stained with Hoechst 33342 (Thermo Fisher Scientific, Cat# H3570). The coverslips were mounted on glass slides using Fluoromount-G (Southern Biotechnology, Cat# 0100-01) with Hoechst 33342. Fluorescence microscopy was performed on a confocal laser microscope (A1RSi, Nikon) and captured with NIS-Elements AR software (Nikon). The area of N-positive cells was quantified using Fiji software implemented in Image J.

### SARS-CoV-2 Reverse Genetics

Recombinant SARS-CoV-2 was generated by circular polymerase extension reaction (CPER) as previously described^39, 43^. In brief, 9 DNA fragments encoding the partial genome of SARS-CoV-2 (strain WK-521, PANGO lineage A; GISAID ID: EPI_ISL_408667)^37^ were prepared by PCR using PrimeSTAR GXL DNA polymerase (Takara, cat# R050A). A linker fragment encoding hepatitis delta virus ribozyme, bovine growth hormone polyA signal and cytomegalovirus promoter was also prepared by PCR. The corresponding SARS-CoV-2 genomic region and the templates and primers of this PCR are summarized in **Extended Data Table 3**. The 10 obtained DNA fragments were mixed and used for CPER^39^. To prepare the GFP-expressing replication-competent recombinant SARS-CoV-2, the fragment 9 in which the *GFP* gene was inserted in the *ORF7a* frame was used instead of the authentic F9 fragment (see **Extended Data Table 3**)^39^.

To produce recombinant SARS-CoV-2, the CPER products were transfected into HEK293-C34 cells using TransIT-LT1 (Takara, cat# MIR2300) according to the manufacturer’s protocol. At one day posttransfection, the culture medium was replaced with Dulbecco’s modified Eagle’s medium (high glucose) (Sigma-Aldrich, cat# R8758-500ML) containing 2% FCS, 1% PS and doxycycline (1 μg/ml; Takara, cat# 1311N). At six days posttransfection, the culture medium was harvested and centrifuged, and the supernatants were collected as the seed virus. To remove the CPER products (i.e., SARS-CoV-2-related DNA), 1 ml of the seed virus was treated with 2 μl TURBO DNase (Thermo Fisher Scientific, cat# AM2238) and incubated at 37°C for 1 h. Complete removal of the CPER products (i.e., SARS-CoV-2-related DNA) from the seed virus was verified by PCR. The working virus was prepared by using the seed virus as described above.

To generate recombinant SARS-CoV-2 mutants, mutations were inserted in fragment 8 (**Extended Data Table 3**) using the GENEART site-directed mutagenesis system (Thermo Fisher Scientific, cat# A13312) according to the manufacturer’s protocol with the following primers: Fragment 8_S D614G forward, 5’-CCA GGT TGC TGT TCT TTA TCA GGG TGT TAA CTG CAC AGA AGT CCC TG-3’; Fragment 8_S D614G reverse, 5’-CAG GGA CTT CTG TGC AGT TAA CAC CCT GAT AAA GAA CAG CAA CCT GG -3’; Fragment 8_S P681R forward, 5’-AGA CTC AGA CTA ATT CTC GTC GGC GGG CAC GTA GTG TA-3’; and Fragment 8_S P681R reverse, 5’-TAC ACT ACG TGC CCG CCG ACG AGA ATT AGT CTG AGT CT-3’, according to the manufacturer’s protocol. Nucleotide sequences were determined by a DNA sequencing service (Fasmac), and the sequence data were analyzed by Sequencher version 5.1 software (Gene Codes Corporation). The CPER for the preparation of SARS-CoV-2 mutants was performed using mutated fragment 8 instead of parental fragment 8. Subsequent experimental procedures correspond to the procedure for parental SARS-CoV-2 preparation (described above). To verify insertion of the mutation in the working viruses, viral RNA was extracted using a QIAamp viral RNA mini kit (Qiagen, cat# 52906) and reverse transcribed using SuperScript III reverse transcriptase (Thermo Fisher Scientific, cat# 18080085) according to the manufacturers’ protocols. DNA fragments including the mutations inserted were obtained by RT-PCR using PrimeSTAR GXL DNA polymerase (Takara, cat# R050A) and the following primers: WK-521 23339-23364 forward, 5’-GGT GGT GTC AGT GTT ATA ACA CCA GG-3’; and WK-521 24089-24114 reverse, 5’-CAA ATG AGG TCT CTA GCA GCA ATA TC-3’. Nucleotide sequences were determined as described above, and sequence chromatograms (Extended Data Fig. 3) were visualized using the web application Tracy (https://www.gear-genomics.com/teal/)^44^.

### Real-Time RT-PCR

Real-time RT-PCR was performed as previously described^43, 45^. Briefly, 5 μl of culture supernatant was mixed with 5 μl of 2 × RNA lysis buffer [2% Triton X-100, 50 mM KCl, 100 mM Tris-HCl (pH 7.4), 40% glycerol, 0.8 U/μl recombinant RNase inhibitor (Takara, cat# 2313B)] and incubated at room temperature for 10 min. RNase-free water (90 μl) was added, and the diluted sample (2.5 μl) was used as the template for real-time RT-PCR performed according to the manufacturer’s protocol using the One Step TB Green PrimeScript PLUS RT-PCR kit (Takara, cat# RR096A) and the following primers: Forward *N*, 5’-AGC CTC TTC TCG TTC CTC ATC AC-3’; and Reverse *N*, 5’-CCG CCA TTG CCA GCC ATT C-3’. The copy number of viral RNA was standardized with a SARS-CoV-2 direct detection RT-qPCR kit (Takara, cat# RC300A). The fluorescent signal was acquired using a QuantStudio 3 Real-Time PCR system (Thermo Fisher Scientific), a CFX Connect Real-Time PCR Detection system (Bio-Rad), an Eco Real-Time PCR System (Illumina) or a 7500 Real Time PCR System (Applied Biosystems).

### Plasmid Construction

A plasmid expressing the SARS-CoV-2 S D614G protein was prepared in our previous study^8^. A plasmid expressing the SARS-CoV-2 S D614G/P681R mutant was generated by site-directed mutagenesis PCR using pC-SARS2-S D614G^8^ as the template and the following primers: P681R Fw, 5’-CCA GAC CAA CAG CCG GAG GAG GGC AAG GTC T-3’ and P681R Rv, 5’-AGA CCT TGC CCT CCT CCG GCT GTT GGT CTG G-3’. The resulting PCR fragment was digested with KpnI and NotI and inserted into the KpnI-NotI site of the pCAGGS vector^46^.

### Pseudovirus Assay

Pseudovirus assay was performed as previously described^8, 43^. Briefly, the pseudoviruses, lentivirus (HIV-1)-based, luciferase-expressing reporter viruses pseudotyped with the SARS-CoV-2 S protein and its derivatives, HEK293T cells (1× 10^6^ cells) were cotransfected with 1 μg of psPAX2-IN/HiBiT^47^, 1 μg of pWPI-Luc2^47^, and 500 ng of plasmids expressing parental S or its derivatives using Lipofectamine 3000 (Thermo Fisher Scientific, Cat# L3000015) or PEI Max (Polysciences, Cat# 24765-1) according to the manufacturer’s protocol. At two days posttransfection, the culture supernatants were harvested, centrifuged. The amount of pseudoviruses prepared was quantified using the HiBiT assay as previously described^8, 47^. The pseudoviruses prepared were stored at –80°C until use. For the experiment, HOS-ACE2 cells and HOS-ACE2/TMPRSS2 cells (10,000 cells/50 μl) were seeded in 96-well plates and infected with 100 μl of the pseudoviruses prepared at 4 different doses. At two days postinfection, the infected cells were lysed with a One-Glo luciferase assay system (Promega, Cat# E6130), and the luminescent signal was measured using a CentroXS3 plate reader (Berthhold Technologies) or GloMax explorer multimode microplate reader 3500 (Promega).

### Western blotting

Western blotting was performed as previously described^48–50^. To quantify the level of the cleaved S2 protein in the cells, the harvested cells were washed and lysed in lysis buffer [25 mM HEPES (pH 7.2), 20% glycerol, 125 mM NaCl, 1% Nonidet P40 substitute (Nalacai Tesque, Cat# 18558-54), protease inhibitor cocktail (Nalacai Tesque, Cat# 03969-21)]. After quantification of total protein by protein assay dye (Bio-Rad, Cat# 5000006), lysates were diluted with 2 × sample buffer [100 mM Tris-HCl (pH 6.8), 4% SDS, 12% β-mercaptoethanol, 20% glycerol, 0.05% bromophenol blue] and boiled for 10 min. Ten microliter of the samples (50 μg of total protein) were subjected to western blotting. To quantify the level of the cleaved S2 protein on virions, 900 μl of the culture medium including the pseudoviruses were layered onto 500 μl of 20% sucrose in PBS and centrifuged at 20,000 × g for 2 h at 4°C. Pelleted virions were resuspended in 1× NuPAGE LDS sample buffer (Thermo Fisher Scientific, Cat# NP0007) containing 2% β-mercaptoethanol, and the lysed virions were subjected to western blotting. For the protein detection, following antibodies were used: mouse anti-SARS-CoV-2 S monoclonal antibody (clone 1A9, GeneTex, Cat# GTX632604), rabbit anti-ACTB monoclonal antibody (clone 13E5, Cell Signaling, Cat# 4970), mouse anti-HIV-1 p24 monoclonal antibody (clone 183-H12-5C, obtained from the HIV Reagent Program, NIH, Cat# ARP-3537), horseradish peroxidase (HRP)-conjugated donkey anti-rabbit IgG polyclonal antibody (Jackson ImmunoResearch, Cat# 711-035-152), and HRP-conjugated donkey anti-mouse IgG polyclonal antibody (Jackson ImmunoResearch, Cat# 715-035-150). Chemiluminescence was detected using SuperSignal West Femto Maximum Sensitivity Substrate (Thermo Fisher Scientific, Cat# 34095) or Western BLoT Ultra Sensitive HRP Substrate (Takara, Cat# T7104A) according to the manufacturers’ instruction. Bands were visualized using the image analyzer, Amersham Imager 600 (GE Healthcare), and the band intensity was quantified using Image Studio Lite (LI-COR Biosciences) or Image J.

### SARS-CoV-2 S-Based Fusion Assay

The SARS-CoV-2 S-based fusion assay was performed as previously described^43^. This assay utilizes a dual split protein (DSP) encoding *Renilla* luciferase (RL) and *GFP* genes, and the respective split proteins, DSP_1-7_ and DSP_8-11_, are expressed in effector and target cells by transfection^49, 51^. Briefly, on day 1, effector cells (i.e., S-expressing cells) and target cells (i.e., ACE2-expressing cells) were prepared at a density of 0.6 to 0.8 × 10^6^ cells in a 6 well plate. To prepare effector cells, HEK293 cells were cotransfected with the expression plasmids for D614G S or D614G/P681R (400 ng) with pDSP_1-7_ (400 ng) using TransIT-LT1 (Takara, Cat# MIR2300). To prepare target cells, HEK293 cells were cotransfected with pC-ACE2 (200 ng) and pDSP_8-11_ (400 ng). In addition to the plasmids above, selected wells of target cells were also cotransfected with pC-TMPRSS2 (40 ng). On day 3 (24 h posttransfection), 16,000 effector cells were detached and reseeded into 96-well black plates (PerkinElmer, Cat# 6005225), and target cells were reseeded at a density of 1,000,000 cells/2 ml/well in 6-well plates. On day 4 (48 h posttransfection), target cells were incubated with EnduRen live cell substrate (Promega, Cat# E6481) for 3 h and then detached, and 32,000 target cells were applied to a 96-well plate with effector cells. RL activity was measured at the indicated time points using a Centro XS3 LB960 (Berthhold Technologies). The S proteins expressed on the surface of effector cells were stained with rabbit anti-SARS-CoV-2 S monoclonal antibody (GeneTex, Cat# GTX635654) and APC-conjugated goat anti-rabbit IgG polyclonal antibody (Jackson ImmunoResearch, Cat# 111-136-144). Normal rabbit IgG (SouthernBiotech, Cat# 0111-01) was used as a negative control. Expression levels of surface S proteins were analyzed using a FACS Canto II (BD Biosciences). RL activity was normalized to the mean fluorescence intensity (MFI) of surface S proteins, and the normalized values are shown as fusion activity.

### Mathematical Modeling for Fusion Velocity Quantification

The following cubic polynomial regression model was employed to fit each of time-series datasets (Fig. 3e):

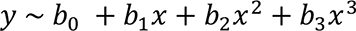

The initial velocity of cell fusion was estimated from the derivative of the fitted cubic curve.

### Neutralisation Assay

Virus neutralisation assay was performed on HOS-ACE2/TMPRSS2 cells using the SARS-CoV-2 S pseudoviruses expressing luciferase (see “Pseudovirus Assay” above). The viral particles pseudotyped with D614G S or D614G/P681R S were incubated with serial dilution of heat-inactivated human serum samples or the RBD-targeting NAbs (clones 8A5, 4A3 and CB6; Elabscience) at 37°C for 1 h. The pseudoviruses without sera and NAbs were also included. Then, the 80 μl mixture of pseudovirus and sera/NAbs was added into HOS-ACE2/TMPRSS2 cells (10,000 cells/50 μl) in a 96-well white plate and the luminescence was measured as described above (see “Pseudovirus Assay” above). 50% neutralisation titre (NT_50_) was calculated using Prism 9 (GraphPad Software).

For the cell-cell fusion neutralisation assay, effector cells of the S-based fusion assay (i.e., S-expressing cells) were incubated with the serially diluted neutralizing antibodies targeting RBD (clones 8A5, 4A3 and CB6; Elabscience) at 37°C for 1 h. Then, target cells were applied and performed the S-based fusion assay as described above (see “SARS-CoV-2 S-Based Fusion Assay” above).

